# Personality predicts dispersal and settlement in a novel habitat – a mesocosm experiment

**DOI:** 10.1101/2024.12.06.627144

**Authors:** J. Gismann, T.G.G Groothuis, F.J. Weissing, M. Nicolaus

**Affiliations:** Groningen Institute for Evolutionary Life Sciences, University of Groningen, The Netherlands

**Keywords:** individuality, three-spined stickleback, RFID, behavioural syndromes, colonisation events

## Abstract

Studying the relationship between dispersal tendencies and personality, or ‘dispersal syndromes’, under ecologically relevant conditions is challenging, especially in fish. Laboratory studies lack environmental complexity and scale, while field-based approaches are often unfeasible. We here mimicked dispersal events; encompassing all three phases of dispersal (departure, transience, settlement) in a large experimental mesocosm containing a gradient of low- and high-quality breeding sites. Using Passive Integrated Transponder (PIT)-tagged three-spined sticklebacks (*Gasterosteus aculeatus*), we remotely monitored the movements and behaviours of two groups, each consisting of 40 individuals of known personality, continuously over two weeks. We tested whether personality variation explained differences in departure timing, transience movements, settlement success and spatial distribution along the environmental gradient. We found evidence for a dispersal syndrome. Specifically, more active/aggressive individuals dispersed quicker in the mesocosm, spent more time on territories and collected more eggs. However, we did not detect phenotype-environment correlations. Our findings highlight the importance of personality-dependent dispersal, with specific behavioural types driving dispersal and settlement success. Furthermore, our mesocosm setup provides unique insights into dispersal dynamics in a system where detailed observations in all dispersal stages are rare. Despite limitations, behavioural experiments in mesocosms serve as a valuable bridge between laboratory and natural systems.

## Introduction

Individual phenotypes play a crucial role during dispersal events, when individuals move to, and ultimately settle in novel habitats. While many studies on dispersal focused on morphological or physiological traits such as body size, body condition or wing length (Bowler & Benton, 2005; Fargallo & López-Rull, 2022), animals primarily interact with their environment through behaviour (Wong & Candolin, 2015). Accordingly, consistent individual differences in behaviours, which are often organised into suites of correlated behaviours (i.e., ’animal personality’), are of major importance during dispersal events (Cote et al., 2011; Mittelbach et al., 2014; Stamps & Groothuis, 2010; Wolf & Weissing, 2012). Such ’personality- dependent dispersal’ or ‘dispersal syndrome’ is a common phenomenon in nature and has repeatedly been observed across many taxa, including birds, mammals, fish, amphibians, and reptiles (Clobert et al., 2009; Cooper et al., 2017; Cote et al., 2011). For example, individuals that showed greater activity levels are often more exploratory, bolder, more aggressive, or less social typically disperse faster and/or farther (Chapple et al., 2012; Cote et al., 2010; Cote & Clobert, 2007; Fraser et al., 2001; Mullon et al., 2017). Dispersal syndromes are often assumed to be adaptive by functionally integrating traits that alleviate the costs of dispersal and/or increase the chances of establishment in novel habitats.

Dispersal can be divided into three consecutive phases: 1) departure, when individuals leave their place of birth (natal dispersal) or breeding (breeding dispersal); 2) transience, when individuals move to and explore novel environments (sometimes referred to as ’floating’); and 3) settlement, when individuals stop moving and settle for, e.g., reproduction (Clobert et al., 2004, 2009; Delgado et al., 2008). Gaining knowledge on the role that individual behavioural differences play in each of these dispersal phases is crucial to better understand the population consequences of dispersal events. For example, studies in western bluebirds (*Sialia mexicana*), showed that new habitats on the edge of the population’s range are exclusively colonised by highly dispersive and aggressive males which outcompete already established mountain bluebirds (*Sialia currucoides*) (Duckworth and Badyaev 2007; Duckworth et al., 2015). Once established, the proportion of aggressive types in the population quickly dropped due to density-dependent performance of the dispersers (Duckworth 2008). This example illustrates that initial settlement success in novel habitats is non-random with respect to behaviour, leading to a spatial clustering of behavioural phenotypes and that dispersal syndromes play a major role in population dynamics.

Studying dispersal syndromes requires quantifying behavioural differences between dispersing and non-dispersing individuals and accurately following the movement of animals over extended periods of time. Through the use of ringing and recent advances in miniaturised GPS tracking technology, researchers are able to decipher animal movement patterns on global scales for e.g., birds, large terrestrial mammals, or sea turtles (Coyne & Godley, 2005; Fancy et al., 1988). Yet, in the underwater environment comparable studies are much more challenging. Studies on aquatic animals are either limited to species that regularly breach the water surface to allow GPS tracking, depend on expensive hardware such as stationary receivers (e.g., acoustic telemetry), or require relatively large tags that may need to be recovered by the researchers (e.g., archival biologgers) (Lennox et al., 2017). As a result, large-scale individual-level data on behaviour and movement of wild fish are hard to obtain. One alternative approach is to sample fish from newly-established populations (i.e., dispersers) and their source populations and to compare behaviours assessed in the lab (e.g., Lukas et al., 2021). While this approach may uncover population behavioural differences, it remains unclear whether and how pre-existing behavioural differences directly contributed to dispersal. Additionally, because fish behaviour is often studied in the laboratory, translating results to natural conditions is challenging. Semi-natural mesocosms, of greater complexity and scale, that allow for detailed behavioural observations, can bridge the gap between small-scale laboratory experiments and field studies. Using such a mesocosm (for details see Ramesh et al., 2023), we here remotely tracked individual movements of large groups of small fish over all phases of dispersal, including settlement and reproduction.

In the mesocosm system (six above-ground interconnected semi-natural ponds), we investigated how consistent differences in individual behaviours relate to dispersal tendencies in the three-spined stickleback (*Gasterosteus aculeatus*). To that end, we performed an experiment with two groups of fish, each consisting of 40 individuals (20 females and 20 males) of known personality (activity and aggression) that we released into a novel environment and monitored continuously over all three phases of dispersal. We focused on activity and aggression as these traits have repeatedly been found to differ between dispersers and non-dispersers (O’Riain et al., 1996; Pocock et al., 2005). By fitting the mesocosm with a limited number of breeding sites that varied along a quality gradient, we monitored male breeding site choice, settlement and mating success. We asked whether and to what extend pre-existing differences in activity and/or aggression contribute to dispersal-related decisions. Specifically, we asked, whether more active/aggressive fish initiate dispersal in the mesocosm quicker during the departure phase; how behavioural differences affect the way individuals sample the environment during the transience phase, and, for males, which individuals are more successful in establishing a territory under competition in the settlement phase. We also explored whether behavioural tendencies predicted the distribution of males over the different types of territory, exploring the potential for phenotype-environment correlations.

## Methods

### Experimental animals

We used three-spined sticklebacks that were born in semi-natural ponds (mesocosms) in summer 2020. These were born from natural mating of wild anadromous sticklebacks caught during inland migration in the northern Netherlands in spring 2020 (Nieuwe Statenzijl: (53°13’54.49’’, 7°12’30.99’’) (Ramesh et al., 2022). The fish were previously used in an experiment investigating the effects of predation cues (olfactory and physical) during development on behaviour as sub-adults (Gismann et al., unpublished data). Since all fish experienced the same treatment during development (3 weeks of predation cues), we are confident that the previous use did not affect the experimental outcomes of the present study. Prior to the experiment, all fish larger than 40mm in total length were tagged with 8mm PIT (‘passive integrated transponder’) tags, allowing individual identification (Trovan, Ltd, Santa Barbara, California) by means of RFID (‘Radio-frequency identification’) technology (for tagging method see Ramesh et al., 2021). In total we ran two experimental rounds on two separate batches with 20 females and 20 males each. One male died over the course of round 2 in the mesocosm, and was subsequently excluded from all analysis (total n= 79 fish).

### Ethical note

Housing of experimental animals, testing of behaviours and PIT tagging were in adherence with the project permit from the Central Committee on Animal Experiments (CCD, the Netherlands; licence number: AVD1050020174084).

### Laboratory tests –Activity and aggression

Fish were housed individually in tanks (15x20 cm) containing a plastic plant for shelter and without visual access to neighbours for ∼ 24h prior to behavioural testing. Housing tanks were filled with a mixture of tap water and water from the mesocosm. For the behavioural trials, fish were carefully transported to test tanks in a water-filled beaker. Four test tanks (same as housing tanks, but lined on all sides and on the bottom with opaque white plastic sheets and filled to a level of 8cm) were placed together in a wooden testing box (60x80x80 cm) to limit disturbances during the trials, illuminated with LED lights. All trials were filmed from the top using Raspberry Pi single-board computers fitted with a camera (Raspberry Pi NoIR Camera Board V2 - 8MP, Raspberry Pi Foundation, UK). Two testing boxes were used simultaneously, such that 8 fish could be tested at a time. After a 5-minute acclimatisation time the behavioural trials began.

‘Activity’ was quantified by the total distance moved (cm) in the empty test tank from 5- minute video recordings using the software ‘EthovisionXT’ (Noldus Information Technology company). Directly thereafter, we remotely lifted an opaque plastic sheet to reveal a mirror on one of the short sides of the arena and recorded the fish’s behaviour for another 5 minutes. We assessed ‘aggression’ by scoring antagonistic behaviours (number of bites and time spent vigorously swimming against the mirror) from video recordings. To avoid observer bias, all videos were scored by the same observer. We noted that fish sometimes interacted with the mirror in non-aggressive ways, such as slow parallel swimming or staying still next to the mirror image, potentially indicating a fish’s willingness to associate with the ’other’ individual. Thus, in contrast to previous studies which simply noted the time spent close to the mirror as a proxy of aggression, our measurements can be readily distinguished from social behaviour.

Every fish was tested twice for both behaviours, with one day in between. For each of the two groups, we tested half of the fish on the same day, such that the laboratory tests took 4 days per group.

### The mesocosm system

The mesocosm consisted of six linearly connected ponds of 1.6m diameter filled to a height of approximately 60cm with water from a nearby waterbody that is similar to natural conditions of wild sticklebacks in the Netherlands. Ponds were connected via tubes (1.5m length; 10cm diameter) and fitted with RDIF antennas at the entrance and exit of each pond to automatically record the fish’s movements between ponds (Fig. 1a). The RFID system was set to a sensitivity of 1 read/s. The reliability of the system was previously evaluated (Ramesh et al., 2023). All ponds contained 2 shelters in the form of artificial plants or plastic tubes.

**Figure 1.**
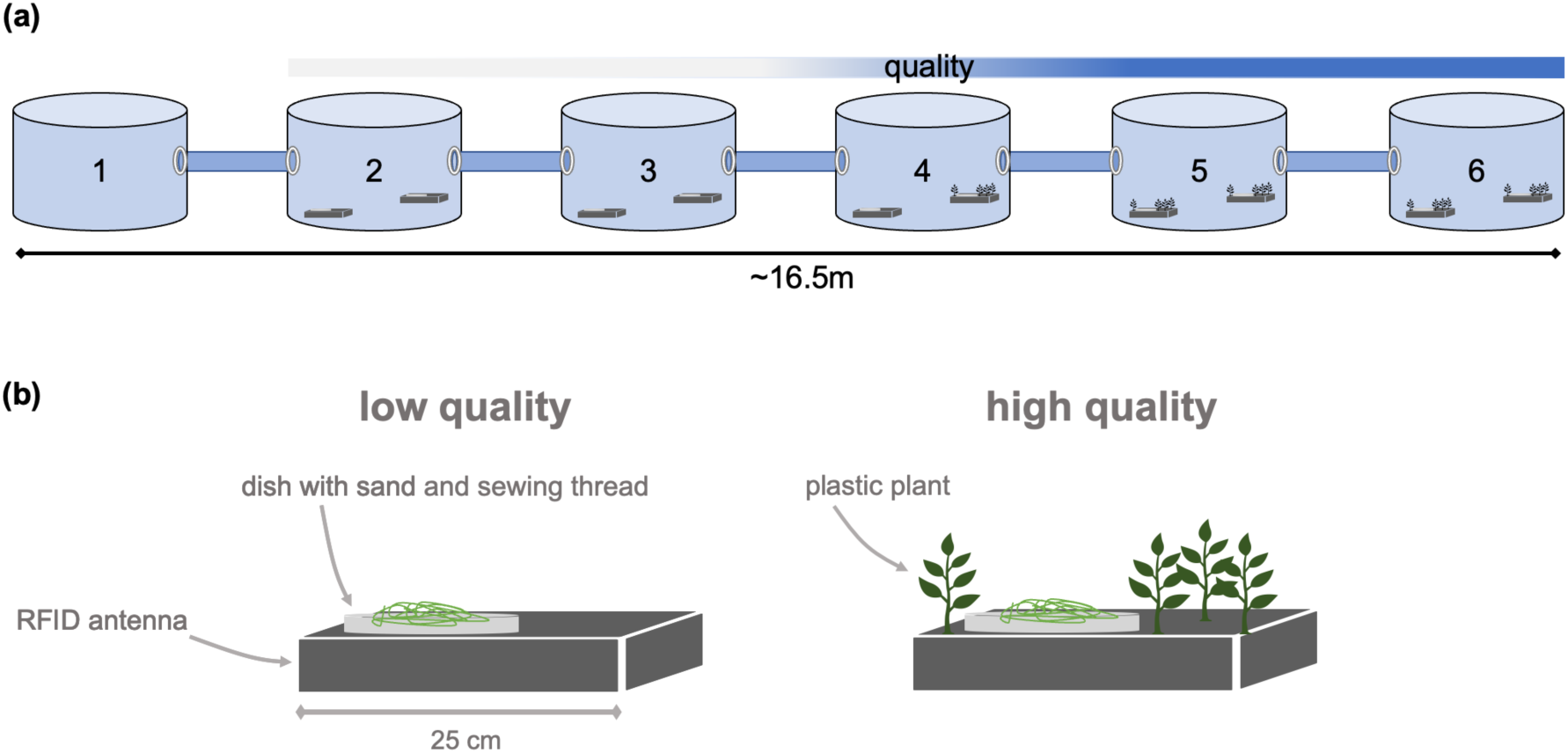
Mesocosm setup and breeding spots. (a) Mesocosm setup containing 2 breeding spots in each pond (except pond 1), with a gradient from low- to high-quality breeding spots toward pond 6. Each pond additionally contained 3 small shelters (not depicted). (b) Low- and high-quality breeding spots placed on top of ’flatbed’ RFID antenna.

Except for the starting pond, each pond contained two breeding spots. These breeding spots consisted of a plastic dish (18cm diameter) filled with 2cm of white aquarium sand (‘lower-quality’) or sand with artificial plants (‘higher-quality’). Male sticklebacks that build nests concealed in plants have previously been shown to exhibit higher mating success (Kraak and Bakker, 1999). We thus expected that breeding spots containing plastic plants would experience higher competition. To aid nest building, all breeding spots in round 2 additionally contained some sewing thread. All breeding spots were placed on top of ’flatbed’ RFID antennas that record the identity of fish that swam within 5 cm of the breeding spot (Fig. 1b). Hence, the presence of territorial or competing males could be identified. We created a gradient from lower- to higher-quality breeding spots the farther fish moved from the starting pond. Ponds 2 and 3 contained two lower-quality breeding spots, pond 4 contained one lower-quality and one higher-quality breeding spot, and ponds 5 and 6 two higher-quality breeding spots each (Fig. 1a).

On the day of testing, fish were released into the first pond of the row, where they were given 4 days in round 1 and 7 days in round 2 to acclimatise. Next, the connection to the other ponds was opened, marking the start of the experiment, and movement and breeding success was monitored for 17 days (10.05.21 - 26.05.21) and 14 days (01.06.21 - 14.06.21) in rounds 1 and 2, respectively. For round 1 we first measured behaviours in the lab and subsequently tested dispersal in the mesocosm. Due to logistic reasons, the order was reversed in round 2.

### Departure phase- Latency to exit the starting pond

From RFID reads we measured each fish’s latency to exit the first pond as the time interval between the start of the experiment (i.e., opening the connector from pond 1 to the other ponds) and reaching pond 2. As all fish exited pond 1, we did not have to define a latency time for fish that never exited the first pond.

### Transience phase- Movements in the mesocosm

From the RFID reads we extracted several measures related to the fish’s behaviour during the transience phase. We extracted movement in the mesocosm as the number of crosses between ponds and crosses between breeding spots within the first 24h after leaving the starting pond (on an individual basis, i.e., based on each individual’s latency to exit the first pond). We chose 24h as the cut-off for the transience phase for several reasons. Firstly, despite its relatively large spatial scale (∼16.5m), the system can be explored rapidly (e.g., on average fish had about 60 pond crosses on the first full day of the experiment in round 1). Secondly, except for 16 individuals, all fish explored the full system (i.e., reached the last pond) within 24h of exiting the first pond. Lastly, we observed that many males started to settle on territories already within the first day of the experiment. Furthermore, as a measure of overall exploration speed, we calculated the *latency to reach the last pond*, as the time it took each fish to reach pond 6 after starting the experiment. Overall, five fish never reached the last pond and were excluded from this aspect of the analysis.

### Settlement- Territory establishment (rounds 1 and 2)

To assess whether and which males successfully established a territory, we applied a number of criteria, based on the reads collected from the antennas underneath the breeding spots. First, as a proxy for the time each male spent on each breeding spot, we counted each male’s total number of daily reads on each breeding spot antenna, (starting from the second day of the experiment). The male with the most reads on a given antenna on a given day was deemed the “occupant” of that specific breeding spot on that day. Because it takes time to establish a territory and to build a nest (Mori, 1993), we considered males as having successfully established a territory if they were the occupant of a breeding spot on three or more consecutive days. We subsequently counted the number of days that each male spent on a territory (only for each males’ first instance of establishing a territory, as we deemed it the more important event regarding male-male competition).

Our criterion of territory occupancy does not distinguish between situations where a single male was the only male to spend time on a given breeding spot. We defined a breeding spot as contested when on the same day two (or more) males had a comparable number of PIT tag reads on that breeding spot. Specifically, we defined a breeding spot as “contested” on a given day if the male with the second-most reads had at least half as many reads as the breeding spot occupant (most read male). In Figure 4, such contested days on a breeding spot are marked by a red cross, as they provide useful information on territory acquisition and conflicts over breeding spots.

### Settlement- Mating success (round 2)

In round 2, we monitored all breeding spots closely for the presence of eggs. We assessed each breeding spot every 2-3 days, by briefly lifting them to the water surface and carefully checking for eggs in nests. When nests were empty, we placed the breeding spot without any further disturbance back on the RFID antenna in the pond. When present, eggs were removed from the nest and counted, and the (partially destroyed) nest was returned to the pond.

### Statistical analyses

#### Laboratory tests

To quantify individual consistency in activity and aggression, we calculated ’raw’ repeatabilities (without the inclusion of fixed effects) using the ’rptR’ package (Stoffel et al. 2017). For activity, we specified a generalised linear mixed model (GLMM) with a Gaussian error distribution and individual ID as a random effect. In the aggression models, we used a Gaussian error distribution for the time spent vigorously swimming and Poisson error distribution with a square root link function for the number of bites and included individual ID as a random effect. We calculated repeatabilities (R) and their 95% confidence intervals (CI) with 1000 bootstraps. For the number of bites, we report ’latent’ (link-scale) approximations (Stoffel et al., 2017). We calculated Pearson correlation coefficients to quantify correlations between behaviours.

#### Personality-dependent dispersal and territory establishment

To test for an association between individual behavioural phenotype and dispersal-related traits, we ran separate linear models for each dispersal behaviour measured in the mesocosm.

We used *mean activity* scores (individual mean over the two trials) as a proxy of personality since activity exhibited the highest repeatability and to avoid collinearity issues (levels of activity and aggression are positively correlated; see results). It is thus important to note that in the rest of the study, more active fish were also more aggressive. At first, all models contained an interaction term between *activity* and *sex* (fitted as a factor with *female* used as reference category). Because the interaction was non-significant and because models without the interaction performed better according to Akaike information criterion (AIC), the interaction term was removed, retaining *sex* as a fixed effect in all final models (except for the male-only model on the days spent on a territory). All models contained the experimental *round* as a fixed effect (with round 1 used as reference category), to control for potential differences between the two batches of fish. For models on the latency to leave the first pond and the latency to reach the last pond (pond 6), Gaussian error distributions were fitted. Data were log-transformed in the model on the latency to reach the last pond. For the number of pond crosses (and breeding spot crosses) during transience and for the number of days on a territory, a negative binomial family was fitted to handle overdispersion, using the glmmTMB package (Brooks et al., 2017) (Table 1). Welch’s two-sample t-tests and Wilcoxon rank-sum tests were performed to test if mean activity level differed between males that acquired a territory or not, acquire eggs or not, and obtained a low quality or a high quality territory. Preliminary analysis revealed that size did not correlate to any lab behaviours (activity and aggression) or dispersal-related traits (latency to leave the first / reach the last pond), and we thus did not include size as a fixed effect in the linear models.

**Table 1.** Model summary of generalised linear models of latency to leave the first pond in hours, hours to reach pond 6 (log-transformed), number of pond crosses during the transience phase, and number of days spent on a territory. Mean activity scores, sex of the fish and experimental round were included as fixed effects with female (F) and Round (1) being used reference categories. Estimates or Incidence Rate Ratios (for neg. binomial model families) are given with their 95% confidence intervals (CI).

| Predictors | Departure Latency |  | Latency to Pond 6 |  | Transience Pond Crosses |  | Days on Territory |  |
| --- | --- | --- | --- | --- | --- | --- | --- | --- |
|  | Estimates | CI | Estimates | CI | Incidence Rate Ratios | CI | Incidence Rate Ratios | CI |
| Intercept | 61.330 *** | 31.583 – 91.076 | 4.152 *** | 3.445 – 4.858 | 147.453 *** | 90.484 – 240.288 | 0.862 | 0.242 – 3.070 |
| Activity | -0.037 * | -0.073 – -0.001 | -0.001 * | -0.002 – -0.000 | 1.000 | 0.999 – 1.000 | 1.002 ** | 1.001 – 1.003 |
| Sex (M) | -11.264 | -29.623 – 7.095 | -0.111 | -0.541 – 0.320 | 0.723 | 0.510 – 1.025 |  |  |
| Round (2) | 12.477 | -5.726 – 30.680 | 0.142 | -0.284 – 0.569 | 0.645 * | 0.454 – 0.917 | 0.611 | 0.237 – 1.574 |
| Observations | 79 |  | 74 |  | 79 |  | 39 |  |

\* $p < 0.05$ \*\* $p < 0.01$ \*\*\* $p < 0.001$

## Results

### Laboratory tests

For activity, we found higher repeatability between trial 1 and trial 2 (R (95% CI) = 0.511 (0.335, 0.67)) than for aggressive behaviours (number of bites: R (95% CI) = 0.393 (0.185, 0.546); time spent vigorously swimming: R (95% CI) = 0.321 (0.104, 0.503)). For subsequent analysis, we calculated individual means in activity, number of bites, and time spent vigorously swimming over the two trials. Both measures of aggression were highly correlated (Pearson ρ = 0.94, p < 0.001; Fig 2a). Activity was positively correlated with both measures of aggression (activity – time spent vigorously swimming: Pearson ρ = 0.47, p < 0.001 (Fig. 2b); activity – bites: Pearson ρ = 0.50, p < 0.001 (Fig. 2c)), indicating the existence of an activity-aggression syndrome. Visual examination of the relationship between activity and aggressive behaviours suggests, that the observed correlations result in part from individuals at both extremes of the activity/aggressiveness distributions. The most active individuals were consistently highly aggressive, and vice versa. Males were overrepresented among the fish with the highest aggression and generally males were more aggressive than females (mean bites males= 80.3; mean bites females= 53.2), while females were less active and overrepresented among the fish with the lowest activity scores (mean activity males= 839.9; mean activity females= 739.9).

**Figure 2.**
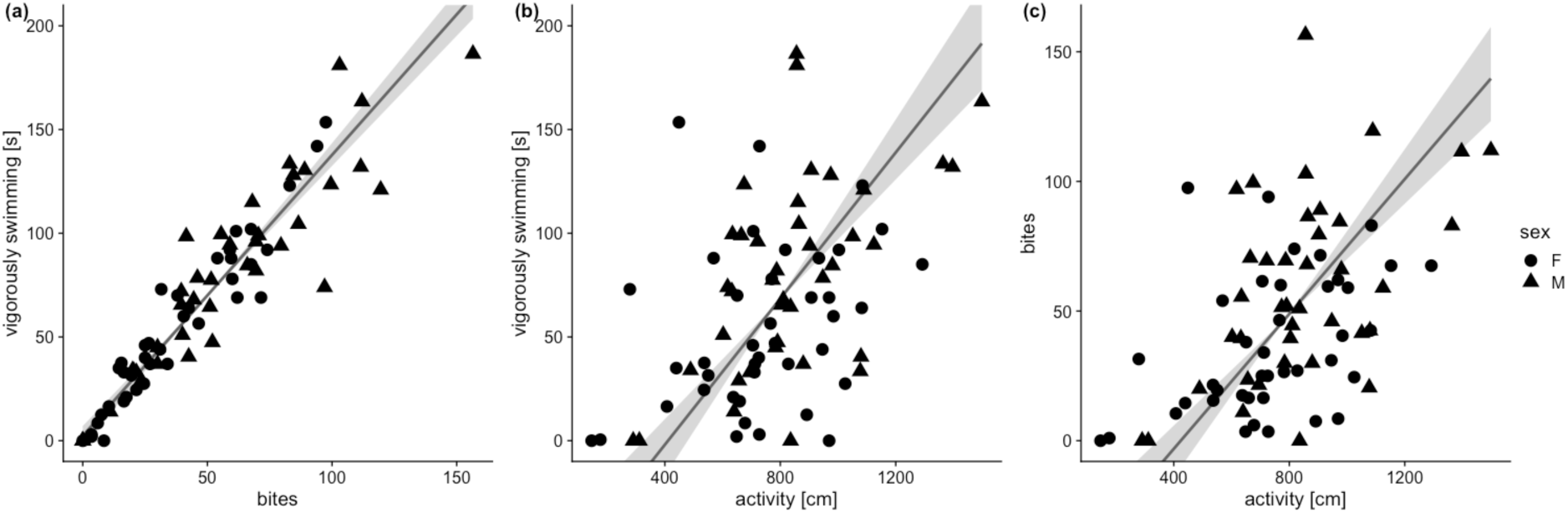
Correlations between behaviours measured in the laboratory. (a) Mean time vigorously swimming plotted against the mean number of bites. (b) Mean time spent vigorously swimming plotted against mean activity (distance moved in cm). (c) Mean number of bites plotted against mean activity. Standardised major axis regression lines are displayed with 95 % confidence intervals (grey shaded area). Females are represented by circles, males by triangles. Sample size for all plots is n= 79.

### Departure

There was considerable variation in the departure time from the first pond, yet all fish in the experiment eventually left the first pond. The fastest fish left within 15 min (0.26h), while the slowest took almost 8 days (188,5h) (median= 18.4h; mean= 32.6h). As indicated by Fig. 3a, more active individuals exited the first pond more quickly than less active ones. The effect was independent of sex and experimental round (Table 1). It is noteworthy, that not all fish fitted to this general pattern. For example, the two fish that displayed the lowest activity levels in the lab, were among the first to leave the first pond. The observed pattern seems to be driven by fish with the high activity scores, which consistently had a low latency to leave the first pond (i.e., variation in latency was greater among less active than among more active individuals). Furthermore, fish that took to the longest to depart were predominantly female.

**Figure 3.**
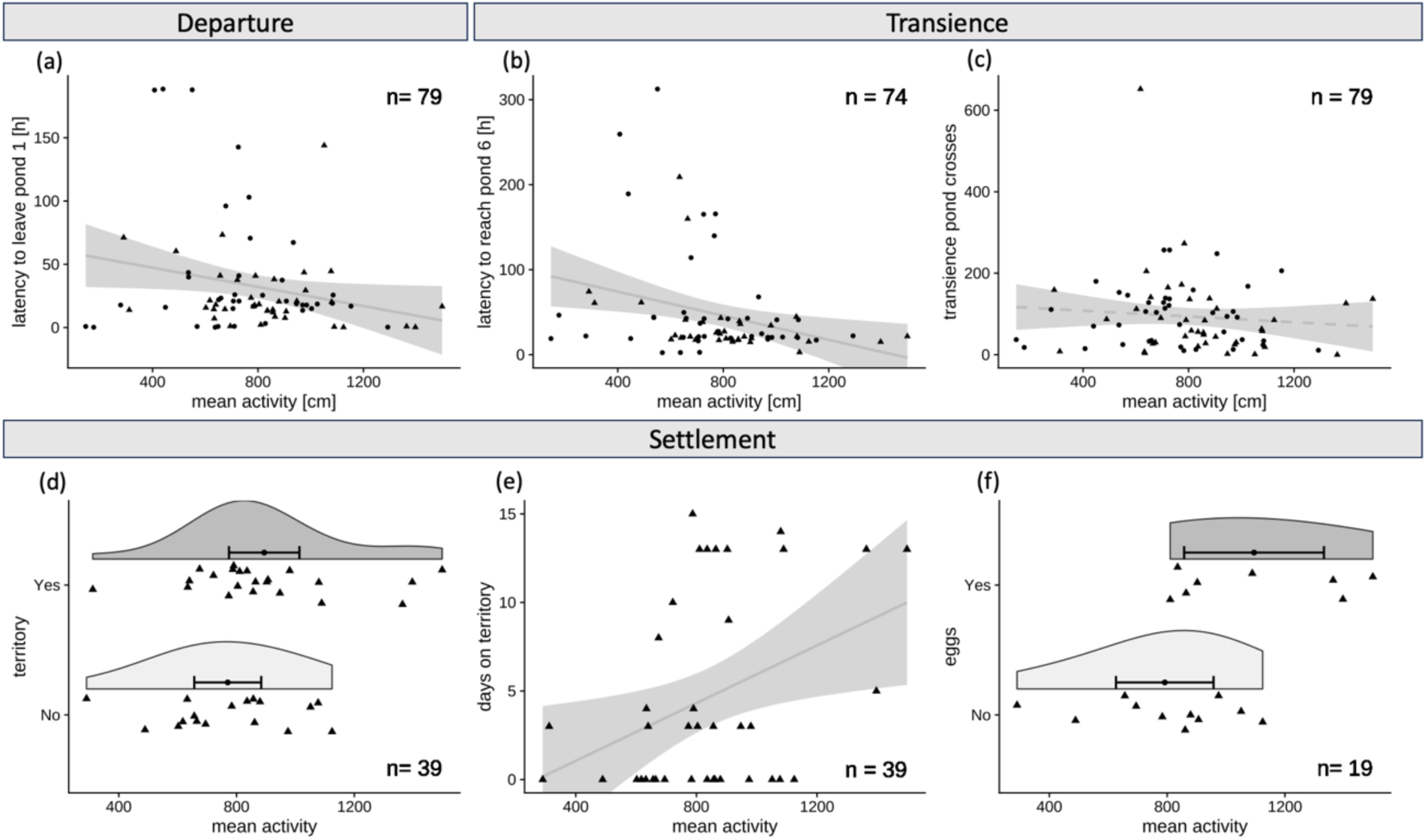
Relationship between activity level and behaviour in the departure, transience, and settlement phase. Departure phase: (**a)** Latency to leave the first pond plotted against mean activity. Transience phase: (b) Latency to reach the last pond plotted against mean activity. **(c)** The number of pond crosses during transience plotted against mean activity. Settlement phase: **(d)** Distribution of mean activity scores of males that successfully acquired a territory (‘Yes’) and males that did not (‘No’). **(e)** The number of days on a territory plotted against mean activity. **(f)** Distribution of mean activity scores of males that acquired eggs (‘Yes’) and males that did not acquire eggs (‘No’) in round 2. Solid lines represent linear regression lines with 95% confidence intervals and dashed lines indicates that the regression coefficient did not differ significantly from zero. In (d) and (f), data points with density kernels are plotted with mean and 95% confidence intervals. Females are represented as circles and males as triangles.

### Transience

All but five fish entered the last pond (Pond 6). The latency to reach the last pond ranged from 2.4h to 312.5h (∼13days) (median= 22.4h; mean= 47.6h). We found that more active fish were faster in reaching the last pond (Fig. 3b; Table 1). The latency to reach the last pond followed a similar distribution as the latency to leave the first pond. Again, variation in latency was greater in less active fish, while the fish with high activity scores were consistently fast to reach the last pond. Again, the fish with the most extreme latency values were mostly female (7 out of 9 fish with latency > 5000 minutes). The number of crosses between ponds (Fig. 3c) and the number of breeding spot crosses (Fig S1a) during the transience phase (the first 24 hours after departure from the first pond) did not covary with activity. In round 2, fish moved less between ponds in the transient phase than fish in round 1 (Fig. S1b; Table 1). No effect of sex was observed in either of the aforementioned models (Table 1).

### Settlement

As presented in Figure 3d, males that acquired a territory exhibited higher levels of activity compared to males that failed to acquire one (mean activity of successful males= 894.0; mean of non-successful males= 770.1), although this trend was not supported by the statistical tests (Welch t-test: t(36.84) = -1.58, p = 0.12; Wilcoxon rank sum test: W= 137, p = 0.16). Yet, considering all fish, more active fish spent more days on territories than less active fish (Fig. 3e; Table 1). In round 2, males that acquired eggs tended to be more active than males that did not (Fig. 3f; Welch t-test: t(13.86) = -2.43, p = 0.03; Wilcoxon rank sum test: W= 22, p = 0.08). The mean activity of males that acquired eggs was 1095.1 (SD = 284.0), vs. 792.0 (SD = 246.6) for those that did not.

Figure 4 provides a more detailed picture of territory establishment in our experiment. For each of the 20 males in round 1 (Fig. 4a) and each of the 19 males in round 2 (Fig. 4c), this figure shows on which experimental day a male was classified as the “owner of a territory”. It also shows for both rounds which of the ten breeding spots were “occupied” or “contested” on each experimental day, and which male was the occupant (Fig. 4b & 4d). In round 1 (Fig. 4a), 13 out of 20 males established a territory, while in round 2 (Fig. 4c), only 9 out of 19 males established a territory. Generally, in round 2, territory establishment happened more quickly and territories were held for a longer period than in round 1. In fact, 7 of the 10 breeding spots had the same occupant throughout the whole experiment in round 2 (Fig. 4d), while breeding spot occupation was much more dynamic in round 1 (Fig. 4c).

**Figure 4.**
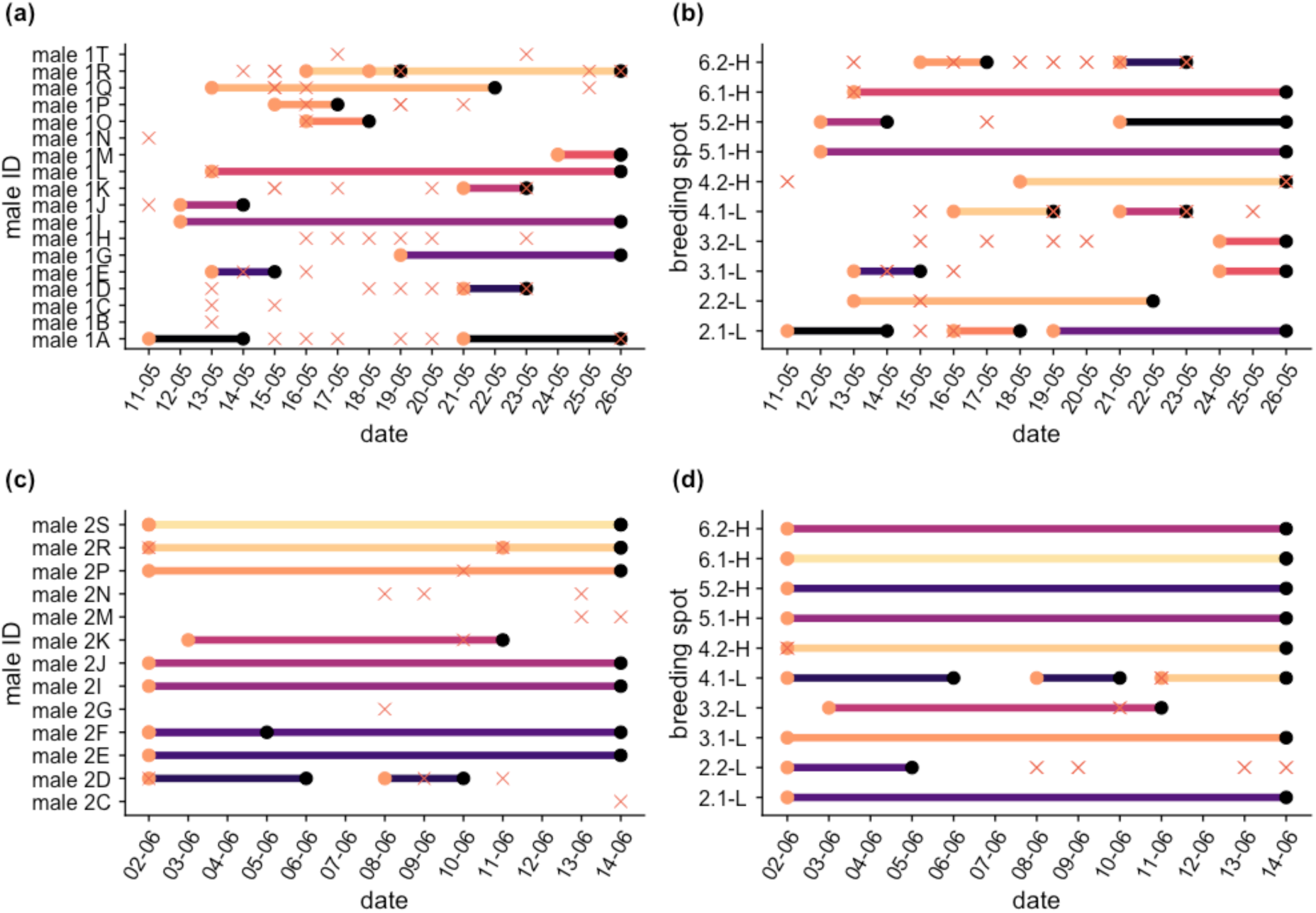
Overview of male territory establishment and breeding spot occupancy. **(a)** Territory establishment for each male that established a territory or had a contested day in round 1. **(b)** Succession of occupancy on each breeding spot in round 1. **(c)** Territory establishment for each male that established a territory or had a contested day in round 2. **(d)** Succession of occupancy on each breeding spot in round 2. Orange dots represent the starting day of territory occupancy and black dots represent the end day. Breeding spot names represent pond number, breeding spot number within pond (e.g., 2.1= pond 2 breeding spot 1) and quality (L= low quality; H=high quality). Red crosses represent contested days on breeding spots. Colours in (b) and (d) correspond to the colours of the males in (a) and (c), respectively.

Some interesting observations can be distilled from the detailed account of breeding spot occupancy in Figure 4. For example, in round 1, *male 1R* first occupied breeding spot *4.1- L* for 4 days and on the third day (18.05.) also began to occupy breeding spot *4.2-H*, the higher-quality breeding spot in the same pond (Fig. 4a & 4b). On the following day, it left the lower- quality breeding spot and remained on the one of higher quality until the end of the experiment. The lower-quality breeding spot was subsequently occupied by *male 1K*. Similarly, in round 2, *male 2F* occupied both breeding spots in pond 2 on the first three days of the experiment (*2.1-L* and *2.2-L*; Fig. 4d). On the subsequent days of the experiment, the male seems to have ‘focused’ on breeding spot *2.1-L*, which it occupied until the end of the experiment. Interestingly, no other male managed to establish a territory on the other, now vacant, breeding spot within the same pond. The fact that breeding spot *2.2-L* was contested on four days in this ‘vacant’ period indicates that several males tried to settle there, but without success.

### Territory quality and spatial distribution of behavioural phenotypes

Over both rounds we found 12 instances of territories established on higher-quality breeding spots and 16 instances of territory establishment on lower-quality breeding spots. On average, males spent more time on higher- than on lower-quality breeding spots (mean time on higher- quality breeding spots: 9.83 days (total of 118 cumulated days); and mean time on lower-quality breeding spots: 5.75 days (total of 92 cumulated days)) (t-test: t(26) = 2.58, p = 0.02; Wilcoxon rank sum test: W = 139.5, p = 0.04), explaining the higher total number of established territories on lower-quality breeding spots.

Breeding spots were arranged such that fish were presented with a gradient from low- to high-quality. However, we did not detect any phenotype-environment correlation (Fig. S2). Males on high- and low-quality breeding spots did not differ in mean activity (mean activity on high-quality = 863.3 vs. mean activity on low-quality = 996.2; t-test: t(26) = -1.18, p = 0.25; Wilcoxon rank sum test: W = 78.5, p = 0.43) but differed however in mean aggression: Males on high-quality breeding spots were less aggressive than males on low-quality breeding spots (mean time spent vigorously swimming on high-quality = 73.3s, vs. low-quality = 111.4s; Welch t-test: t(20.99) = -2.38, p = 0.03; Wilcoxon rank sum test: W = 47.5, p = 0.03; Fig S3).

## Discussion

This study aimed to investigate how individual differences in activity and aggression contribute to dispersal-related choices and outcomes over the different phases of dispersal in three- spined sticklebacks. As previously reported in this species (Bell & Stamps, 2004; Dingemanse et al., 2007), our study confirms that more active fish were more aggressive. Importantly, and as expected, we further found evidence for personality-based dispersal in all three phases of dispersal. Compared to less active-aggressive individuals, more active-aggressive fish departed quicker and moved faster during the transience phase, although behavioural types did not differ in the way they sampled the environment. Personality did not predict territory acquisition but, among successful settlers, more active-aggressive males overall spent more time on a territory, and, in the second round, acquired more eggs. Territory quality did not affect the spatial distribution of personality types, thereby not supporting the existence of phenotype-environment correlations along the manipulated gradient. We discuss below the implications of our findings.

We show that high levels of activity (and aggression) were positively associated with faster dispersal and higher mating success. Uncertainty around the mechanisms underlying such behavioural correlations often limits our understanding of the eco-evolutionary implications of dispersal syndromes (Nicolaus et al., 2022). For example, behavioural differences between dispersers and residents might rather be the consequence than the cause of dispersal if dispersers adjust their behaviour to conditions experienced during or after dispersal (Holekamp, 1986; Hoset et al., 2011). Here we tested aggression and activity shortly before the dispersal experiment in round 1, and shortly after the dispersal experiment in round 2. Because the same relationships were found between activity/aggression and faster dispersal across both experimental rounds, it is unlikely that the dispersal syndrome results from short-term plasticity. Instead, our study highlights that in our population, pre-existing personality variation led to a subset of the population driving dispersal. Should our results extend to stickleback dispersal in natural settings, we hypothesise that post-glacial repeated colonisation events of freshwater habitats by marine sticklebacks (Bell & Foster, 1994; McKinnon & Rundle, 2002) may have been driven by a subset of the ancestral population exhibiting a ‘dispersive’ behavioural phenotype (high activity and aggression levels). Newly established populations in sticklebacks may thus differ in the composition of behavioural phenotypes compared to their source population, at least initially. Since dispersal related traits often have a genetic basis, dispersal syndromes can result in founder effects which have important consequences for subsequent adaptation in novel environments and may ultimately play a role in speciation (Santos et al., 2012; Via, 1999).

We further found, that more active and more aggressive phenotypes were not only more dispersive but also more likely to collect eggs during reproduction under competition. As a result, the proportion of more active-aggressive individuals could increase in the next generations of the newly established population. Invasive cane toads (*R. marina*) in Australia provide an example of such a phenomenon. Individuals at the colonisation front have been found to be more exploratory and risk-prone than individuals from the core-populations (Gruber et al., 2017a). Differential selection likely favoured dispersal-related traits that aid in the search for food and shelter. Because dispersing toads are more likely to breed among each other and because these traits have a genetic basis (Gruber et al., 2017b), the frequency of exploratory/risk-prone phenotypes increased over time, accelerating the evolution of dispersal-enhancing traits (Cayuela et al., 2020).

One intriguing question arising from our results is thus why not all individuals in the population display these beneficial dispersal-related behaviours. Several mechanisms can explain the persistence of behavioural heterogeneity within newly established populations. Firstly, dispersal may be costly. Predation pressure, for example, may be stronger on more dispersive individuals (Wolf & Weissing, 2012). Secondly, selection on dispersal syndromes likely fluctuates and/or is context-dependent. For example, in pied flycatchers (*Ficedula hypoleuca*), the strength of the association between dispersal and aggression and its associated fitness differed between years, which highlights that specific behavioural phenotypes are favoured in some years but not in others (Nicolaus et al., 2022). Similarly, in western bluebirds (*Sialia mexicana*) the fitness of highly aggressive dispersing individuals was shown to be density-dependent (Duckworth 2008). Additionally, the structure of behavioural syndromes can be less stable than typically assumed. A long-term study on collared flycatchers (*Ficedula albicollis*) demonstrated that the strength and direction of behavioural correlations generally differed between the years (Garamszegi et al., 2015). Accordingly, the most active- aggressive-dispersive individuals in our experiment may thus only be favoured during the initial phase of colonisation/settlement and will likely be subjected to heterogeneous selection over time. Other behavioural types may be favoured at a later point in time when, for example, population densities change.

Our results did not reveal personality-related differences in sampling of the environment during the transience phase. Following theories on coping styles, one could have expected that highly active, aggressive and dispersive individuals (‘proactive’), would explore their environment quickly but superficially, whereas ‘reactive’ individuals would explore slowly and thoroughly (Koolhaas et al., 1999; Réale et al., 2010). The lack of personality-related differences in sampling could be an artefact of the timing of the experiment, which took place relatively late in the sticklebacks’ reproductive period. The fact that all but one breeding spot was occupied from the first day of the experiment in round 2 highlights that males attempted to reproduce as soon as the opportunity arose, without prolonged sampling. The reproductive imperative may have masked individual variation in exploration. Additionally, mesocosm complexity may have been insufficient to demand thorough exploration.

Against our expectation, we did not detect a clear phenotype-environment correlation along the environmental gradient: male aggression but not activity level differed between high- or low-quality breeding spots and, while all breeding sites were ultimately occupied, males spent more time on high-quality ones. Thus, opposed to previous studies (Bensky & Bell, 2022; Pearish et al., 2013), we can here not conclusively discern whether different personality types exhibited preferences (or outcompeted others) for specific habitats. In three-spine sticklebacks, it was shown that personality types consistently used different microhabitats in the wild based on diet (Pearish et al. 2013). Such phenotype-environment correlations can lead to feedback loops which affect levels of phenotypic variation within a population and can be an important source of variation in fitness (Wolf & Weissing, 2010). Additionally, if individuals select habitats that match their phenotype (‘matching-habitat choice’; Edelaar et al., 2008); selection will be relatively weak compared to individuals that are excluded from their preferred habitats due to e.g., social competition. The absence of phenotype-environment correlations in our experiment could thus lie in our choice of the environmental gradient. It is plausible that the pressure for timely reproduction and competition for breeding opportunities were so large, that males simply did not discriminate between breeding sites.

Altogether, our findings add to the growing literature on dispersal syndromes, with personality traits being integrated with dispersal tendencies and associated with differential fitness outcomes. We present rare evidence of such syndrome in a small fish where individual- level observations in the wild are limited. Using an experimental mesocosm, we observed dispersal behaviours across all three phases for up to 17 days. Fast dispersers experienced greater mating success, indicating potential positive feedback loops. While no clear phenotype-environment correlations were observed, they could exist along other gradients like social environments. Further mesocosm experiments exploring phenotype assortment and distribution promise exciting avenues. For fish, mesocosms offer a viable alternative to unfeasible studies in the wild, though they do not fully capture important ecological pressures that counteract dispersal advantages, such as predation, energetic limitations, or heterospecific competition. More long-term work quantifying fluctuating selection pressures is thus warranted given population’s large behavioural variation.

## Supplementary material

**Fig. S1:**
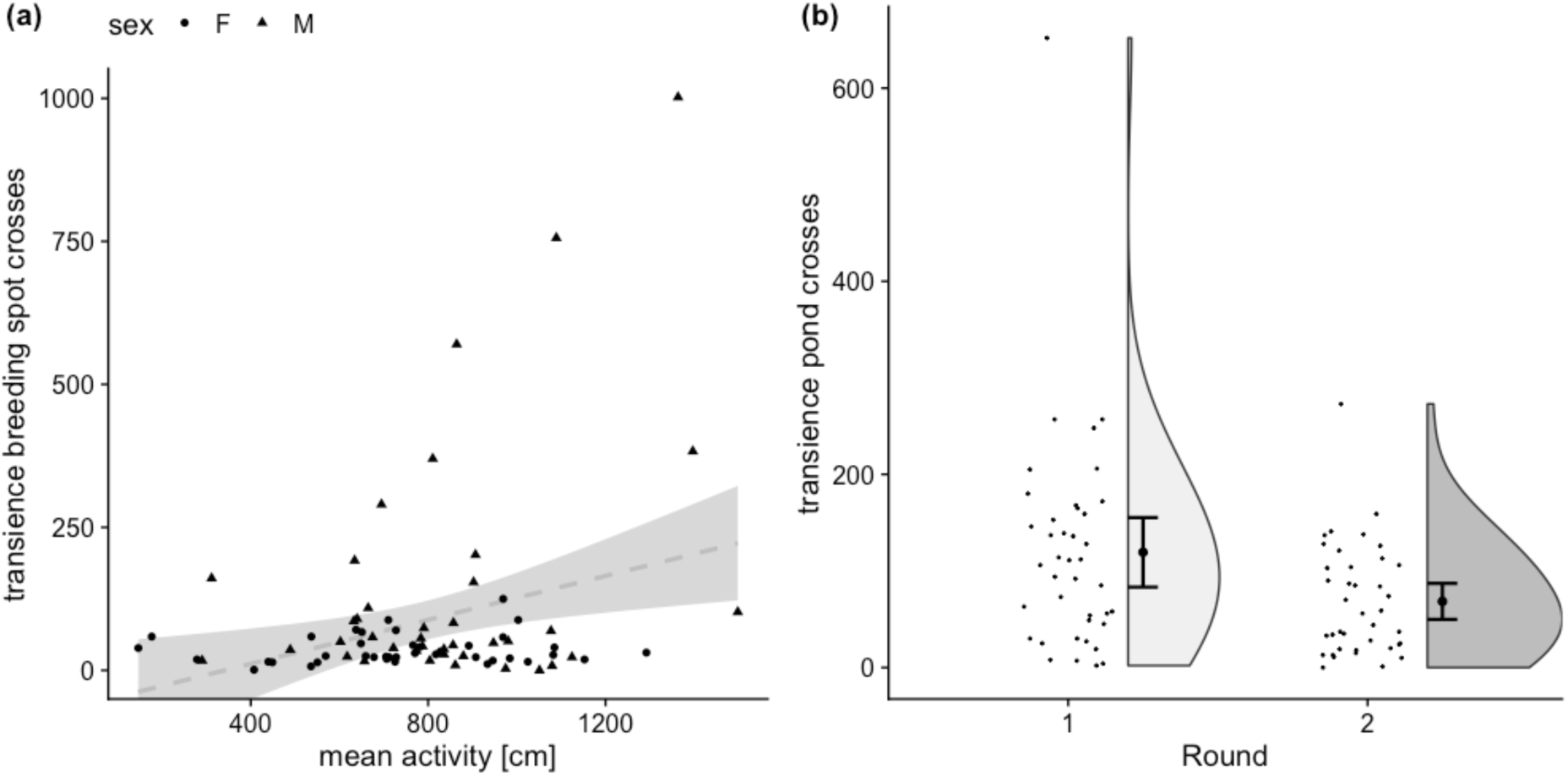
Additional plots on transience behaviours. **(a)** Number of crosses between breeding spots in the first 24h after departure plotted against mean activity. We did not find that more active fish moved more often between breeding spots than less active fish. The dashed grey line (with 95% confidence interval) indicates that the regression coefficient did not significantly differ from zero. Females are represented as circles and males as triangles. **(b)** The number of pond crosses during the transience phase plotted by round, showed that fish in round 1 crossed more often than in round 2. Data points with density kernels are plotted with mean and 95% confidence intervals.

**Fig. S2:**
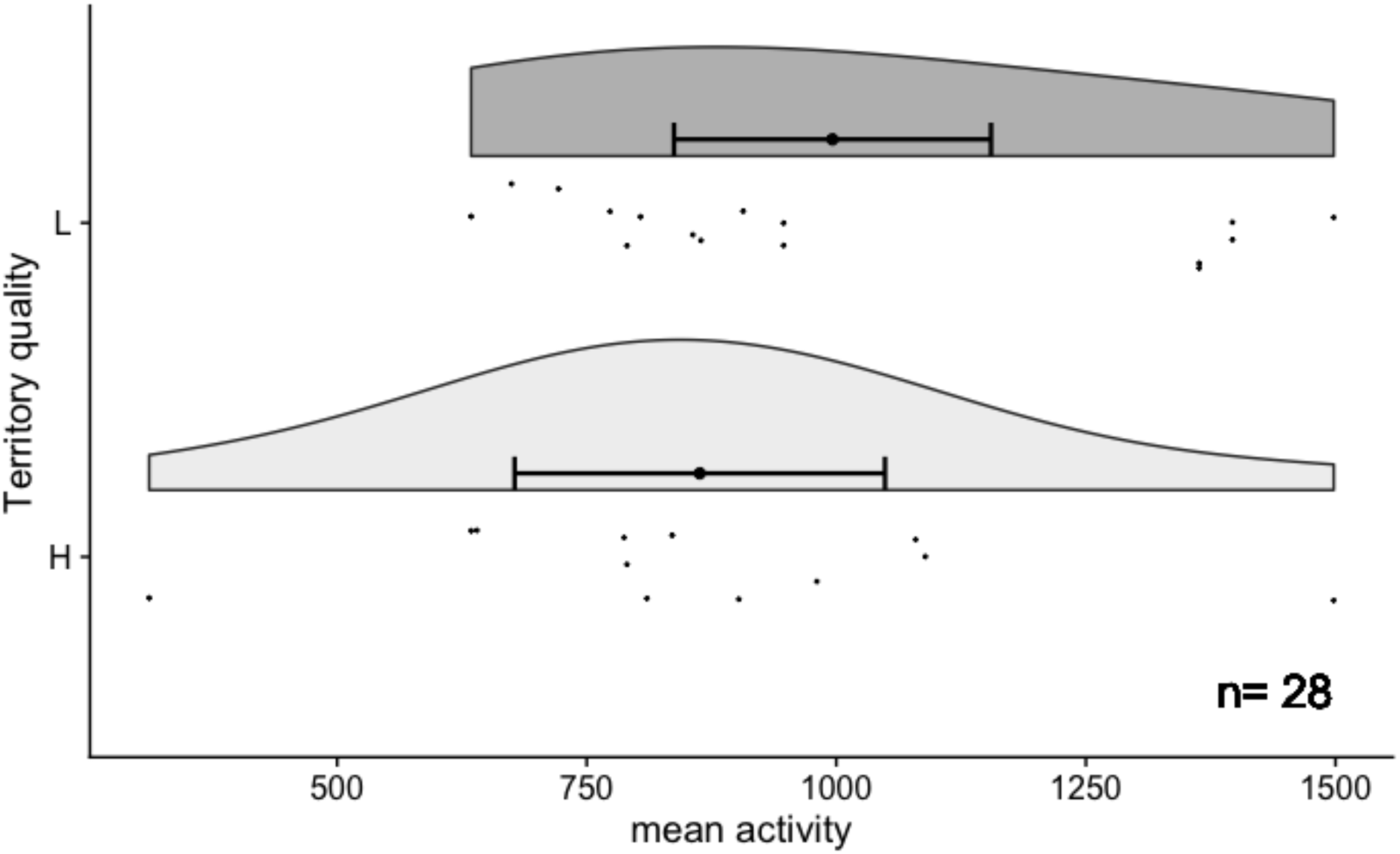
Relationship between male activity and territory quality. L= lower-; H= higher-quality territories. Data points with density kernels are plotted with mean and 95% confidence intervals. Males on higher-quality breeding spots had a mean activity score of 863.3, and males on lower-quality breeding spots had a mean activity score of 996.2 (t-test: t(26) = -1.18, p = 0.25; Wilcoxon rank sum test: W = 78.5, p = 0.43). In total we found 28 instances of territory establishment.

**Fig. S3:**
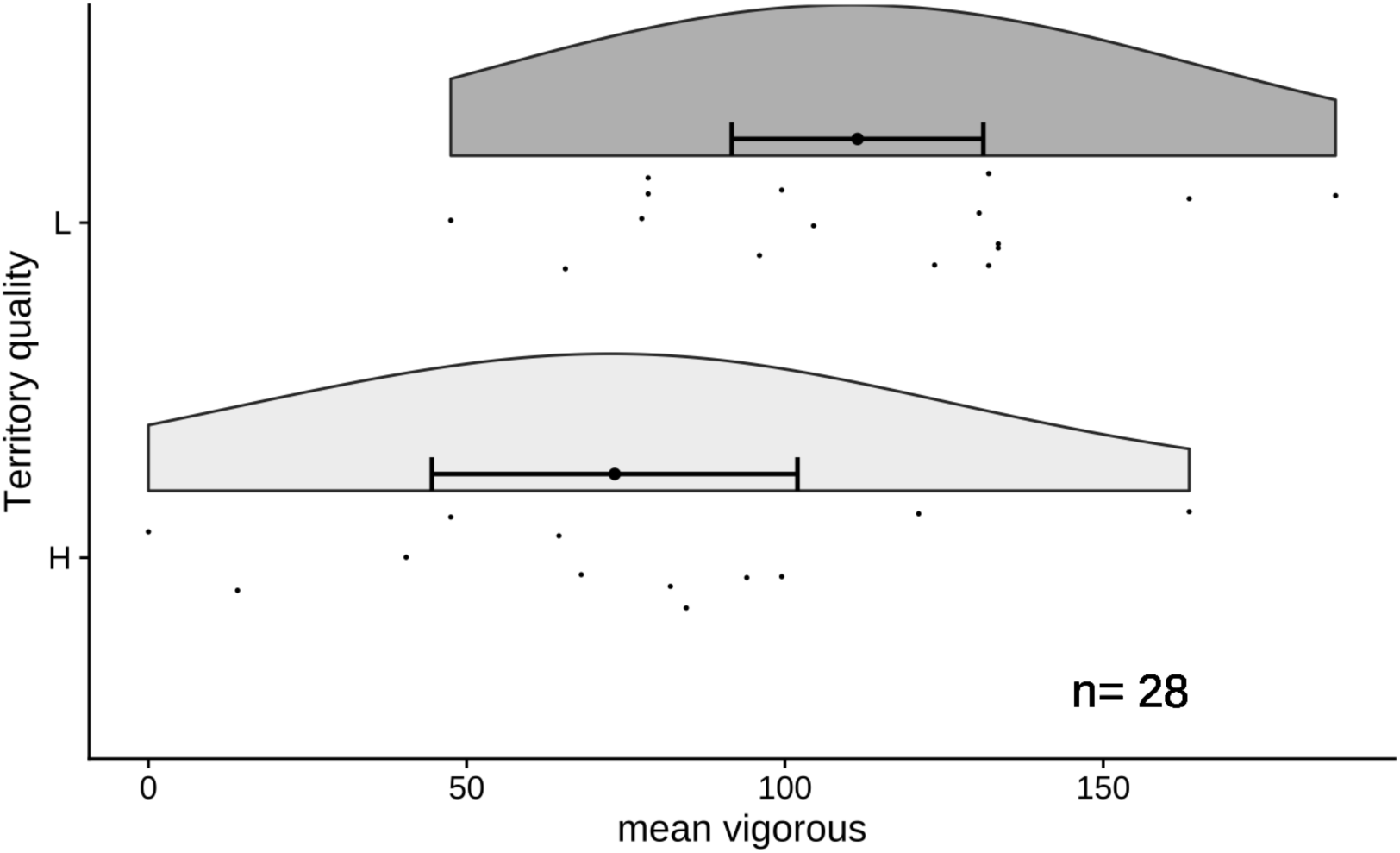
Relationship between male aggression and territory quality. L= lower-; H= higher-quality territories. Data points with density kernels are plotted with mean and 95% confidence intervals. We found that male son lower-quality territories were more aggressive than males on higher-quality territories. Males that occupied higher-quality breeding spots had a mean time spent vigorously swimming of 73.3s, while males that occupied lower-quality breeding spots had a mean time spent vigorously swimming of 111.4s) (Welch t-test: t(20.99) = -2.38, p = 0.03; Wilcoxon rank sum test: W = 47.5, p = 0.03). In total we found 28 instances of territory establishment.

**Table S1:**
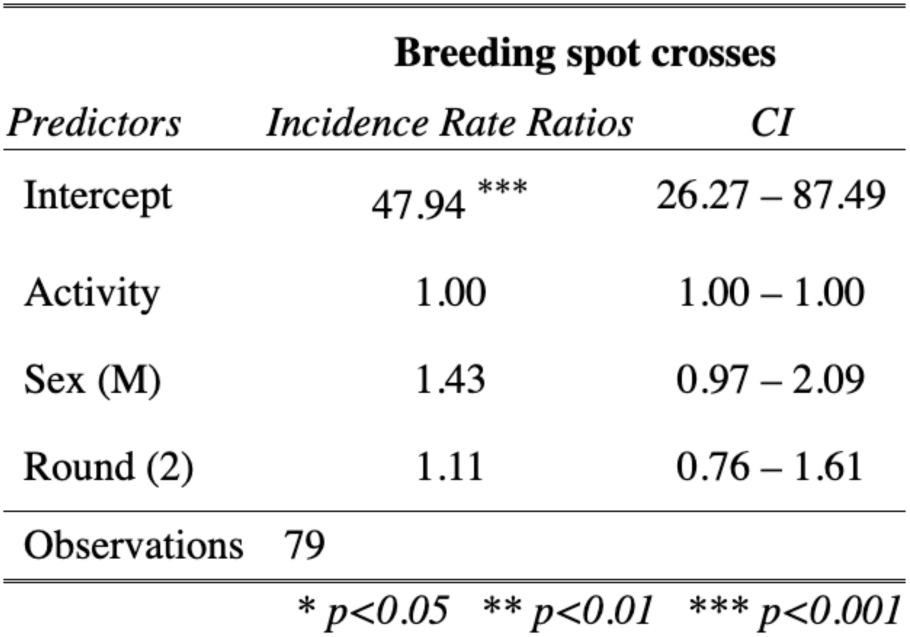
Results of linear regression analyses for the dependence of the number of crosses between breeding spots during the transience phase on mean activity scores. We used generalised linear mixed models including the sex of the fish and the experimental round as fixed effects. Incidence Rate Ratio for neg. binomial model family are given with their 95% confidence intervals. Sex (F) and Round (1) are the reference categories.

